# Proteome, bioinformatic and functional analyses reveal a distinct and conserved metabolic pathway for bile salt degradation in the *Sphingomonadaceae*

**DOI:** 10.1101/2021.05.19.444901

**Authors:** Franziska M. Feller, Lars Wöhlbrand, Johannes Holert, Vanessa Schnaars, Lea Elsner, William W. Mohn, Ralf Rabus, Bodo Philipp

**Author notes:** Corresponding author: Bodo Philipp; Adress: Institut für Molekulare Mikrobiologie und Biotechnologie, Westfälische Wilhelms-Universität Münster, Corrensstr. 3, 48149 Münster, Germany. Current address: Medizinisches Labor Wahl, Lüdenscheid, Germany (Lea Elsner).

## Abstract

Bile salts are amphiphilic steroids with a C_5_ carboxylic side chain with digestive functions in vertebrates. Upon excretion, they are degraded by environmental bacteria. Degradation of the bile-salt steroid skeleton resembles the well-studied pathway for other steroids like testosterone, while specific differences occur during side-chain degradation and the initiating transformations of the steroid skeleton. Of the latter, two variants via either Δ^1,4^- or Δ^4,6^-3-ketostructures of the steroid skeleton exist for 7-hydroxy bile salts. While the Δ^1,4^- variant is well-known from many model organisms, the Δ^4,6^-variant involving a 7-hydroxysteroid dehydratase as key enzyme has not been systematically studied. Here, combined proteomic, bioinformatic and functional analyses of the Δ^4,6^-variant in *Sphingobium* sp. strain Chol11 were performed. They revealed a degradation of the steroid rings similar to the Δ^1,4^-variant except for the elimination of the 7-OH as key difference. In contrast, differential production of the respective proteins revealed a putative gene cluster for side-chain degradation encoding a CoA-ligase, an acyl-CoA dehydrogenase, a Rieske monooxygenase, and an amidase, but lacking most canonical genes known from other steroid-degrading bacteria. Bioinformatic analyses predicted the Δ^4,6^-variant to be widespread among the *Sphingomonadaceae*, which was verified for three type strains which also have the predicted side-chain degradation cluster. A second amidase in the side-chain degradation gene cluster of strain Chol11 was shown to cleave conjugated bile salts while having low similarity to known bile-salt hydrolases. This study signifies members of the *Sphingomonadaceae* remarkably well-adapted to the utilization of bile salts via a partially distinct metabolic pathway.

**Importance:** This study highlights the biochemical diversity of bacterial degradation of steroid compounds, in particular bile salts. Furthermore, it substantiates and advances knowledge of a variant pathway for degradation of steroids by sphingomonads, a group of environmental bacteria that are well-known for their broad metabolic capabilities. Biodegradation of bile salts is a critical process due to the high input of these compounds from manure into agricultural soils and wastewater treatment plants. In addition, these results may also be relevant for the biotechnological production of bile salts or other steroid compounds with pharmaceutical functions.

## Introduction

Bile salts are multifunctional steroidal compounds that act as detergents in the digestion of lipophilic nutrients and exhibit signaling function in vertebrates (1, 2). The amphiphilic character of mammalian bile salts is determined by a carboxylic C_5_ side chain at C17 and one to three hydroxy groups on the steroid nucleus. Bile salts are produced from cholesterol in the liver, conjugated to taurine or glycine via amide bonds and excreted into the gastrointestinal tract. In the intestine, free bile salts are released by deconjugation catalyzed by bile-salt hydrolases produced by intestinal bacteria (3). Although most bile salts are reabsorbed (4, 5), about 0.4 – 0.6 g of bile salts are excreted per day by each human (6), adding up to about 18 t of bile salts excreted per year by the population of a city with 100,000 inhabitants.

Upon excretion, bile salts become an energy and carbon source for environmental bacteria (7, 8) and several bile-salt degrading bacteria have been isolated from soils and aquatic habitats (9–12). These include *Rhodococcus jostii* RHA1 (13), *Comamonas testosteroni* CNB-2 and TA441 (14), *Pseudomonas stutzeri* Chol1 (9), *Pseudomonas* sp. strain DOC21 (11), *Azoarcus* sp. strain Aa7 (12), and *Sphingobium* sp. strain Chol11, formerly *Novosphingobium* sp. strain Chol11 (10). Aerobic bile salt degradation proceeds similar to the degradation of other steroids such as cholesterol and can be divided into different phases (Fig 1) (7, 8, 15): 1) Oxidation of the A-ring, 2) side-chain degradation, 3) oxygenolytic cleavage of ring B, and 4) oxygenolytic and hydrolytic degradation of the remaining *seco-*steroid. The first three steps may occur simultaneously (16–18).

**Fig 1.**
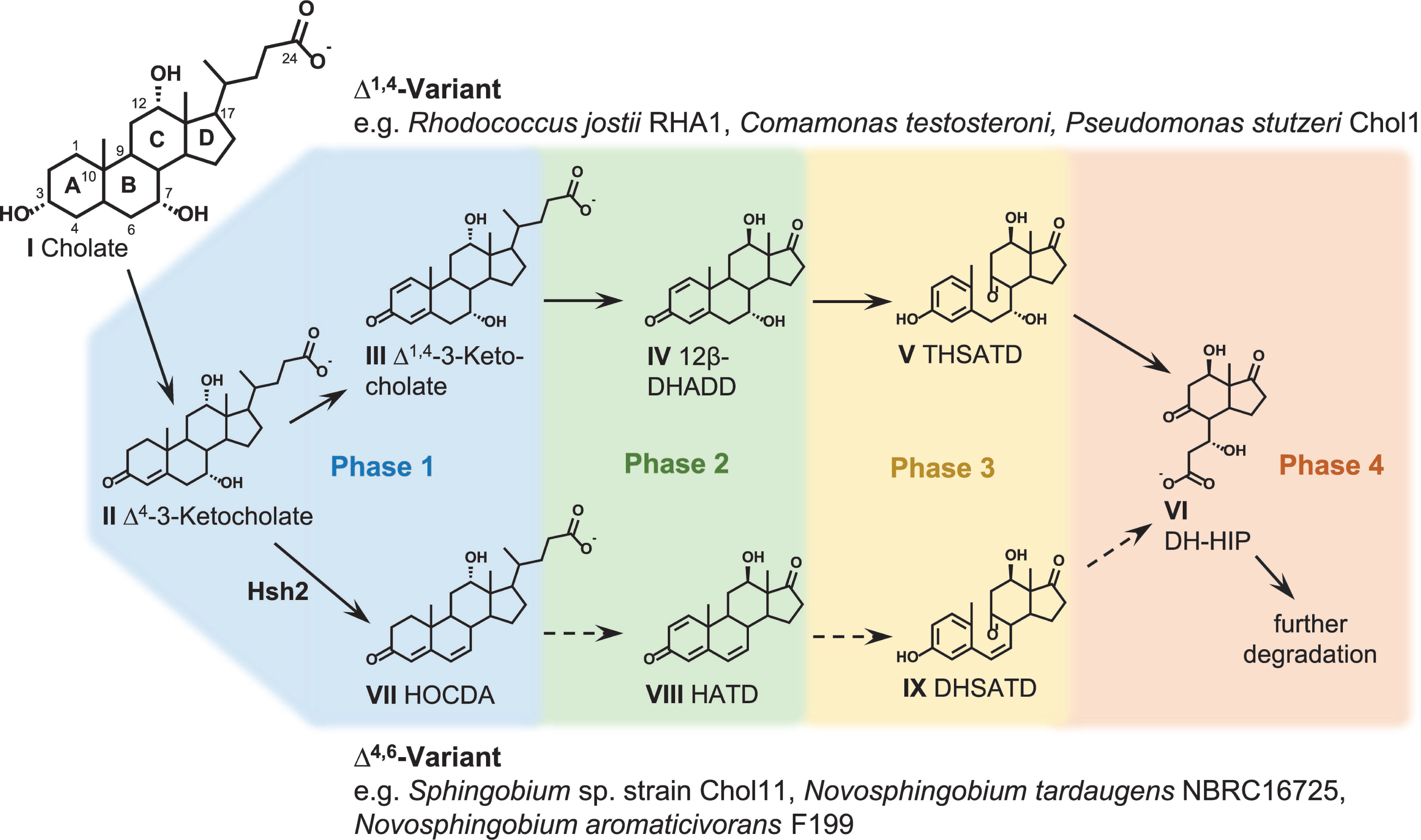
Section of cholate degradation via the Δ^1,4^- and Δ^4,6^-variants of the 9,10-*seco*- pathway. Solid lines: known reactions, dotted lines: predicted reactions. Blue: A-ring oxidation and 7-OH elimination (phase 1), green: side-chain degradation (phase 2), yellow: B-ring cleavage (phase 3), orange: degradation of the 9,10-*seco-*steroid (phase 4). Compound abbreviations: IV, 12β-DHADD (7α,12β-dihydroxy-androsta-1,4-diene- 3,17-dione); V, THSATD (3,7,12-trihydroxy-9,10-*seco*-androsta-1,3,5(10)-triene-9,17- dione); VI, DH-HIP (3’,7-dihydroxy-H-methyl-hexahydro-indanone-propanoate); VII, HOCDA (12α-hydroxy-3-oxo-4,6-choldienoic acid); VIII, HATD (12-hydroxy-androsta- 1,4,6-triene-3,17-dione); IX, DHSATD (3,12β-dihydroxy-9,10-*seco*-androsta-1,3,5(10),6-tetraene-9,17-dione).

In *R. jostii* RHA1, *C. testosteroni* TA441, and *P. stutzeri* Chol1, bile-salt degradation proceeds through the well-elucidated 9,10-*seco* pathway. In phase 1, oxidative reactions at the A-ring generate intermediates with a Δ^1,4^-3-keto structure of the steroid skeleton (7, 13, 14). During degradation of the trihydroxy bile-salt model-substrate cholate (I in Fig 1), this leads to formation of Δ^1,4^-3-ketocholate (III) (16). In phase 2, the bile-salt side-chain is degraded by the successive release of acetyl-CoA and propionyl-CoA (16, 19, 20). In actinobacteria such as *R. jostii* RHA1 acetyl-CoA is predicted to be released by β-oxidation (13). In proteobacteria such as *P. stutzeri* Chol1 acetyl-CoA is released by an aldolase-mediated cleavage reaction and subsequent oxidation of the resulting aldehyde group (19, 21). Both mechanisms of side-chain degradation result in intermediates with C_3_ carboxylic side chains (16, 17, 21, 22). In actino- as well as proteobacteria, this C_3_ side chain is released as propionyl-CoA by a second cycle of aldolytic cleavage reactions (22–24) resulting in C_19_ steroids, so-called androsta-1,4- diene-3,17-diones (ADDs); in the case of cholate, 7,12β-dihydroxy-ADD (12β-DHADD, IV) is formed (9, 13, 20).

In phase 3, degradation of the steroidal ring system is initiated by the introduction of a hydroxy group at C9 by the monooxygenase KshAB (17), which leads to spontaneous opening of the B-ring driven by the aromatization of ring A. This produces 9,10-*seco* intermediates such as 3,7,12-trihydroxy-9,10-*seco*-androsta-1,3,5-triene-9,17-dione (THSATD, V). Phase 4 starts with the *meta*-cleavage of the aromatic A-ring and hydrolytic cleavage of the former A-ring, which results in *H*-methyl-hexahydro-indanone- propanoate (HIP, VI) intermediates (7, 8). At this stage of degradation, intermediates from differently hydroxylated bile salts are channeled into a common pathway in *P. stutzeri* Chol1 (23). In this process, the former 12-OH is removed and a hydroxy group at former C7 is introduced into 7-deoxy-bile salt derivatives during β-oxidation of the former B-ring. Further degradation of HIPs proceeds via β-oxidation of the former B-ring and hydrolytic cleavages of rings C and D (25, 26).

In contrast to this well elucidated pathway, degradation of 7-hydroxy-bile salts such as cholate (I in Fig 1) proceeds differently in *Sphingobium* sp. strain Chol11 (10), but can also be divided into the four phases. After the initial formation of Δ^4^-3-keto-intermediates such as Δ^4^-3-ketocholate (II in Fig 1) in phase 1, the hydroxy group at C7 is eliminated by the 7α-hydroxy steroid dehydratase Hsh2 (27). This leads to the formation of a double bond in the B-ring and to Δ^4,6^-intermediates such as 12-hydroxy-3-oxo-chol-4,6- dienoate (HOCDA, VII) (10). This variant of the pathway will be referred to as Δ^4,6^- variant, in contrast to the Δ^1,4^-variant described above (28). As Δ^4,6^-derivatives of ADDs such as 12-hydroxy-androsta-1,4,6-triene-3,17-dione (HATD, VIII) can be found in culture supernatants of strain Chol11 growing with cholate (10), side chain degradation seems to be the next phase (phase 2) of degradation. This is initiated by CoA-activation catalyzed by CoA-ligase SclA (29). In contrast to the model organisms using the Δ^1,4^- variant, *Sphingobium* sp. strain Chol11 growing with cholate produces no intermediates with a shortened side chain that can be found in culture supernatants (10). Interestingly, many genes for side-chain degradation known from other model organisms are missing in strain Chol11 (29, 30). The fact that several KshA homologs and many HIP degradation proteins are encoded in the genome of strain Chol11 suggests that phases 3 and 4, cleavage of the B-ring and further degradation of the *seco*-steroid, proceed similar to the 9,10-*seco*-pathway (29).

To further elucidate bile-salt degradation in strain Chol11 via the Δ^4,6^-pathway variant, differential proteome analyses of substrate-adapted cells were performed. For this, cholate (I in Fig 1) was used as a model substrate for the Δ^4,6^-variant. This was compared to the 7-deoxy bile salt deoxycholate (XX in Fig 7), since 7-deoxy bile salts cannot be degraded via Δ^4,6^-intermediates (27), and therefore require either completely different pathways or variations of one common pathway. As a further reference substrate, 12β-DHADD (IV in Fig 1) was used, because it does not possess a side chain and therefore might reveal proteins that are specific for side-chain degradation.

## Results and Discussion

### Proteome analyses reveal three gene clusters encoding bile-salt degradation

To assess inducibility of cholate degradation in *Sphingobium* sp. strain Chol11, cholate- and glucose-grown cells were compared. Suspensions of cholate-adapted cells depleted cholate more quickly than those adapted to glucose (Fig S1A). When protein synthesis was inhibited by chloramphenicol, cholate was still completely degraded by cholate- adapted cells, but only a low percentage was depleted by glucose-adapted cells paralled by continuous production of HOCDA (VII in Fig 1) (Fig S1A+B). These findings indicate inducibility of cholate degradation in strain Chol11, with enzymes for A-ring oxidation and 7α-dehydroxylation constitutively produced but those for degradation of the side chain and the steroid nucleus requiring *de novo* synthesis in glucose-grown cells.

Thus, the proteomic profiles of cholate-, deoxycholate-, and 12βDHADD-grown cells were compared to that of glucose-grown cells using two-dimensional difference gel electrophoresis (2D DIGE), whole-cell shotgun proteomics, and analyses of the membrane protein enriched fractions (Tab S1). In total, 44.6% of the 3,550 predicted proteins of strain Chol11 were detected (Fig S2A). Proteins from the COG categories, “inorganic ion transport and metabolism” and “lipid transport and metabolism” showed the highest increase in abundance in cholate-grown cells compared to glucose-grown cells (Fig S2B).

The majority of proteins detected with significantly higher abundances in steroid- versus glucose-grown cells are encoded on chromosome 2 where most predicted steroid-degradation genes in strain Chol11 are located (30). 75 proteins had significantly higher abundances in cells grown on bile salts versus glucose, according to 2D-DIGE analysis.

Most of these proteins are encoded in three gene clusters (Fig 2), of which one (cluster 3) was previously predicted to encode steroid-degradation (29).

**Fig 2.**
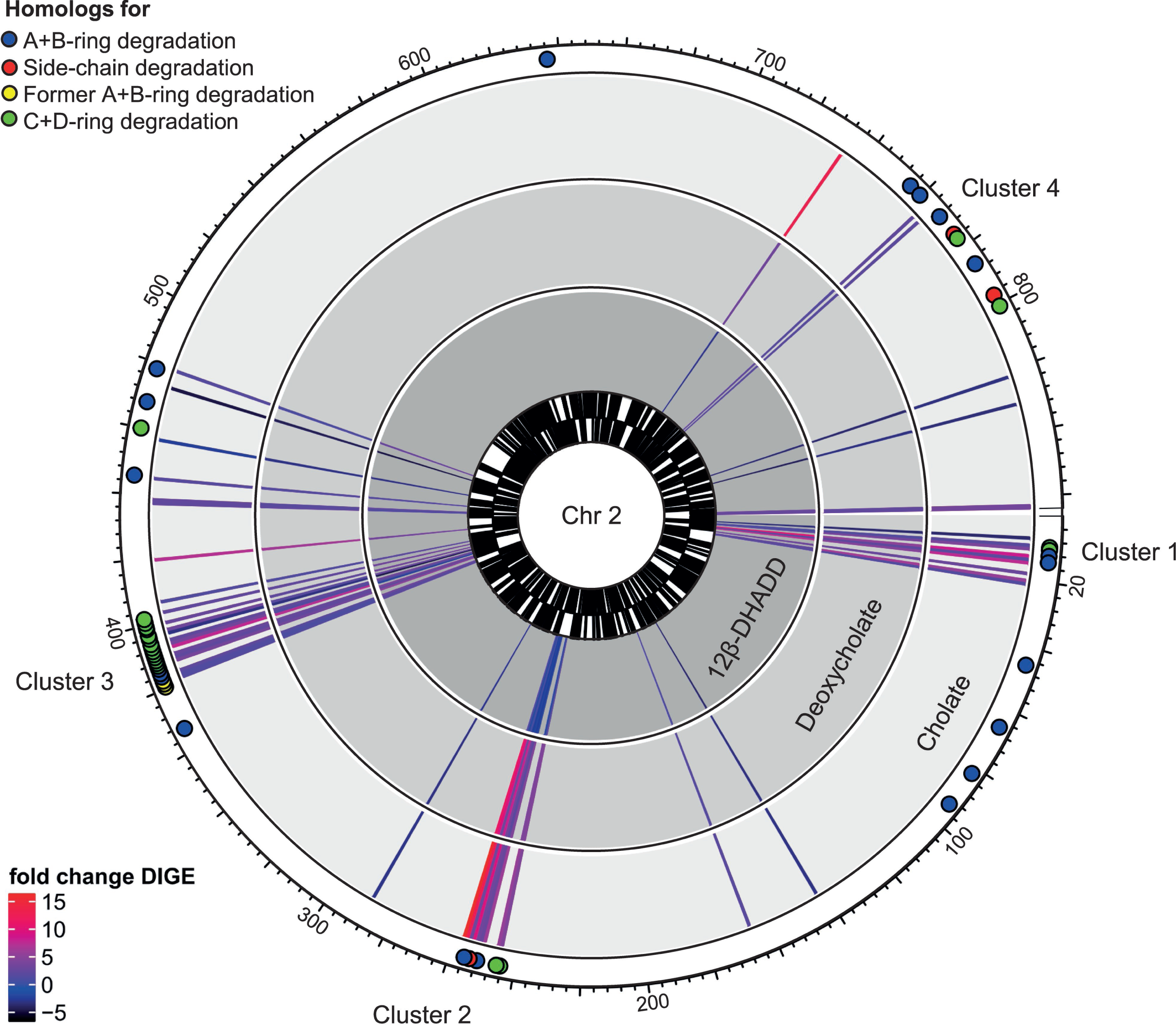
Chromosome 2 of *Sphingobium* sp. strain Chol11 highlighting steroid-specific proteome signatures. Distinct steroid degradation gene clusters are labelled and numbered according to the text. Rings from outside to inside: (i) Localization of genes encoding orthologs to enzymes involved in different phases of steroid degradation based on reciprocal BLASTp analyses similar to (29) (Table S1), (ii-iv) Genes encoding proteins with >|1.5|-fold abundance changes under at least one test condition according to 2D-DIGE with glucose-grown cells as reference, abundance changes for cells grown with (ii) cholate, (iii) deoxycholate or (iv) 12β-DHADD, (v) coding sequences transcribed in clockwise direction and (vi) coding sequences transcribed in counter-clockwise direction.

The finding, that the same set of proteins is produced in higher quantities during growth with both bile salts and most proteins were also produced for 12β-DHADD degradation indicates that degradation of these steroids generally involves the same proteins. This implies that both 7-hydroxy- and 7-deoxy-bile salts are degraded via the same pathway. Notably, however, a subset of proteins encoded in gene cluster 2 is differentially abundant, depending on the presence of a side chain. This implicates the proteins encoded in gene cluster 2 might be specific for side-chain degradation. Based on proteome and bioinformatic analyses, we compiled a model of bile-salt degradation in strain Chol11 (Fig 3).

**Fig 3.**
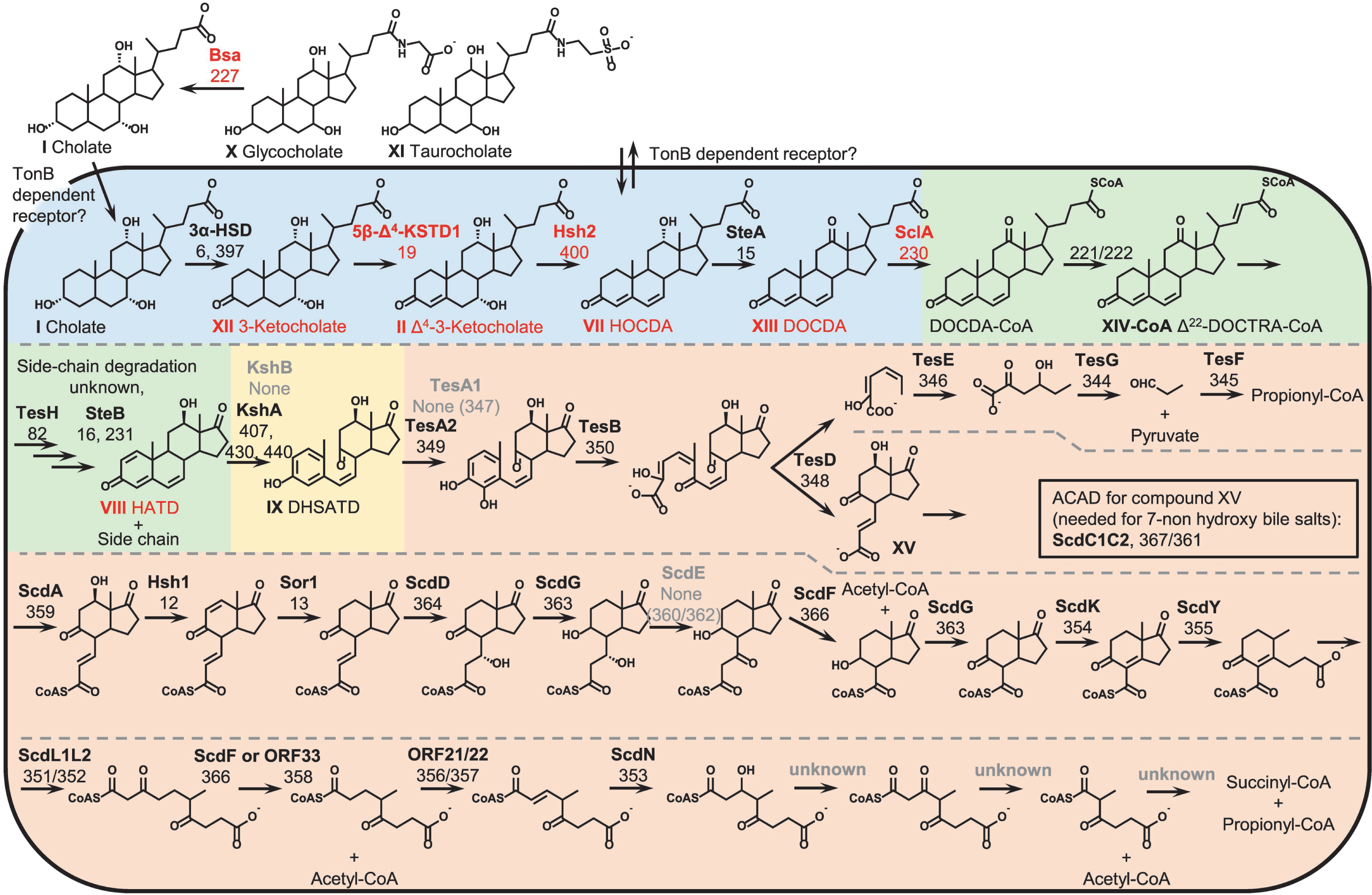
Proposed model of bile-salt degradation in *Sphingobium* sp. strain Chol11 based on integrated bioinformatic, differential proteome and physiological analyses. Gene locus tags are shortened to their numbers, e.g. 6 for Nov2c6. Color coding: (i) enzymes: red, function demonstrated experimentally; black, function predicted due to reciprocal BLASTp analyses with characterized steroid degradation proteins; grey, function predicted due to chromosomal location and automatic annotation; (ii) intermediates: red, intermediates identified in cell-free supernatants of strain Chol11 cultures degrading cholate; black, intermediates predicted due to pathway prediction; (iii) background: blue, A-ring oxidation and 7-OH elimination (phase 1); green, side-chain degradation (phase 2); yellow, B-ring cleavage (phase 3); orange, degradation of the 9,10-*seco*-steroid (phase 4). Compound abbreviations: XIII, DOCDA (3,12-dioxo-4,6-choldienoc acid); XIV, Δ^22^-DOCTRA (3,17-dioxo-4,6,(22*E*)-choltrienoic acid); XV, Δ^3^-7-OH-HIP (7-hydroxy-H-methyl-hexahydro-indanone-3-propenoate).

### Steroid transport

Several TonB-dependent receptor proteins (COG category “inorganic ion transport and metabolism”) were among the membrane proteins with the increases in abundance, including Nov2c232 (gene cluster 2), Nov2c378 (near gene cluster 3), and Nov2c659 (near gene cluster 4). TonB-dependent outer membrane transporter proteins (called TonB-dependent receptor protein) require the accessory proteins TonB, ExbB, and ExbD in the inner membrane to form a functional TonB-system (31). These accessory proteins were identified in the membrane protein-enriched fractions of cells grown on all substrates with similar Mascot scores, indicating constitutive formation (Nov1c1853- 1856). Transporters for the uptake of bile salts are not known in Proteobacteria but there are indications that TonB-dependent receptors could be involved. First, they are generally known to participate in the import of complex growth substrates such as lignin degradation compounds (32). Second, in *Novosphingobium tardaugens* NBRC16725, a TonB-dependent receptor was upregulated in estradiol-grown cells, and a corresponding deletion mutant showed reduced growth with estradiol (33).

The TonB-dependent receptor Nov2c232 (28 % identity to the TonB-dependent receptor that was deleted in *N. tardaugens* NBRC16725) is encoded in the putative side-chain degradation cluster in close vicinity to a gene encoding a transporter of the Major Facilitator Superfamily (MFS) (Nov2c225), and both were more abundant in cholate- and deoxycholate-grown cells (Fig 4, Table S1). These could be involved in transport of bile salts as well as early degradation intermediates such as Δ^4^-3-ketocholate (II in Fig 1) or HOCDA (VII), that are found in strain Chol11 culture supernatants (10, 28). Such transient extracellular accumulation of intermediates (10) is a common for bile-salt degrading Proteobacteria and has been observed during bile-salt degradation in soil (34). Thus, the above transporters could alternatively be involved in intermediate efflux.

**Fig 4.**
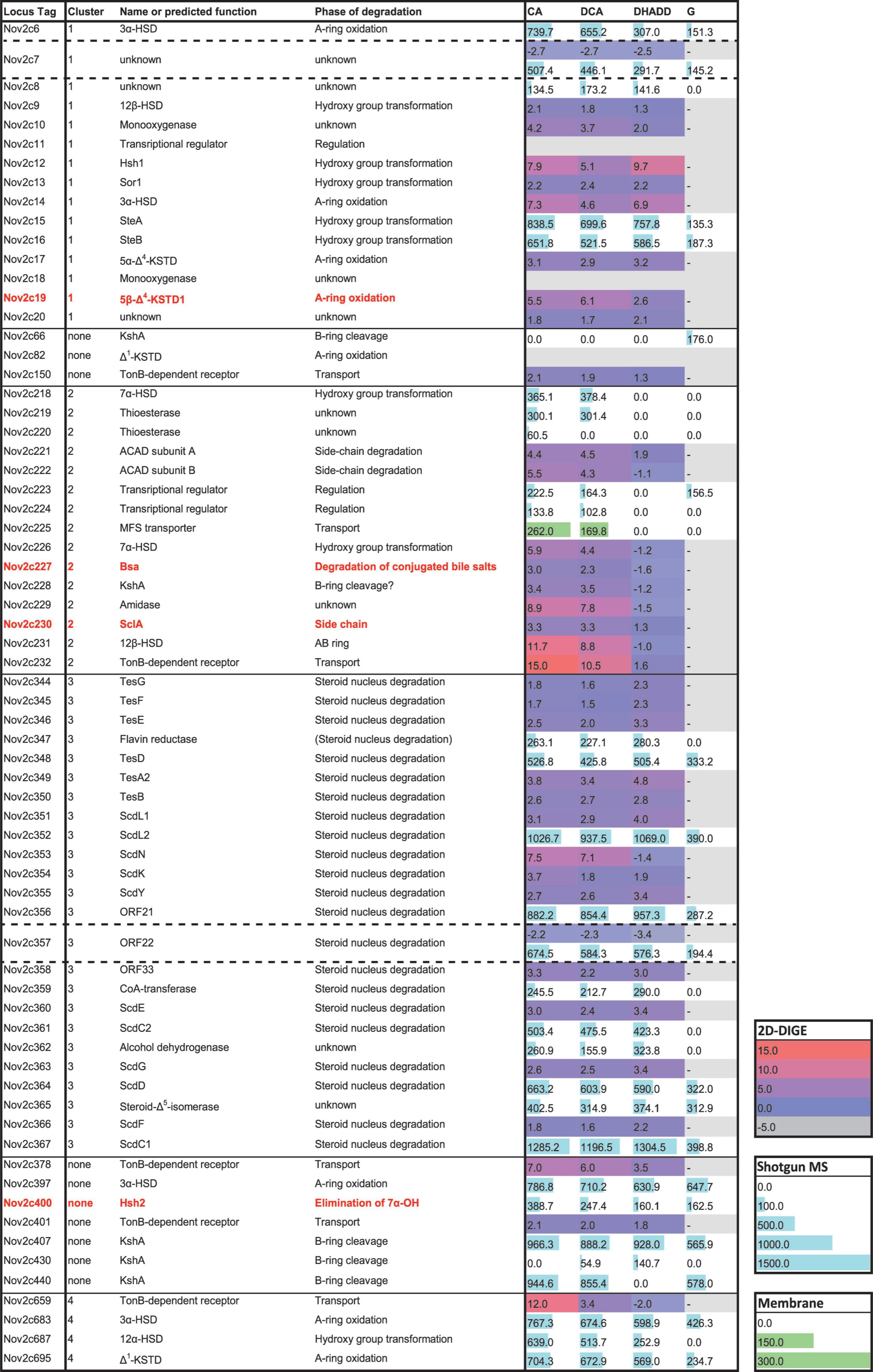
Differential protein expression in *Sphingobium* sp. strain Chol11 adapted to growth with cholate (CA, I in Fig 1), deoxycholate (DCA, XX in Fig 7), and 12β-DHADD (DHADD, IV in Fig 1) compared to glucose (G) as reference state. Proteins are ordered according to their location on chromosome 2. In most cases, results from only one protein identification and quantitation approach are displayed: 2D-DIGE (black to red gradient: protein abundance fold changes with glucose-grown cells as reference state), shotgun-MS analysis (Mascot scores in blue) or MS analysis of the membrane protein- enriched fraction (Mascot scores in green). In case of protein identification by multiple approaches, priority was for 2D DIGE followed by shotgun analyses. Proteins not identified with any method are shown in grey. The complete dataset can be found in Table S1. Red font: known proteins with experimentally verified functions.

### Phase 1: A-ring oxidation and B-ring dehydratation (blue section in Fig 3) 3α-Hydroxysteroid dehydrogenase

The first step of bile-salt degradation is the oxidation of 3-OH to a keto group by a 3α-hydroxy steroid dehydrogenase (3α-HSD) (27, 35). A putative 3α-HSD (Nov2c6) is encoded in gene cluster 1 (Figs 4,5). This protein was present in lower abundances in glucose- versus steroid-grown cells (Fig 4). Additional putative 3α-HSDs, Nov2c397 and Nov2c683, are encoded close to gene clusters 3 and 4, respectively, and were not differentially expressed. This as well as Nov2c6 being present in low abundances in glucose-adapted cells is consistent with biotransformation experiments showing that oxidation of the 3-OH is constitutive in strain Chol11.

**Fig 5.**
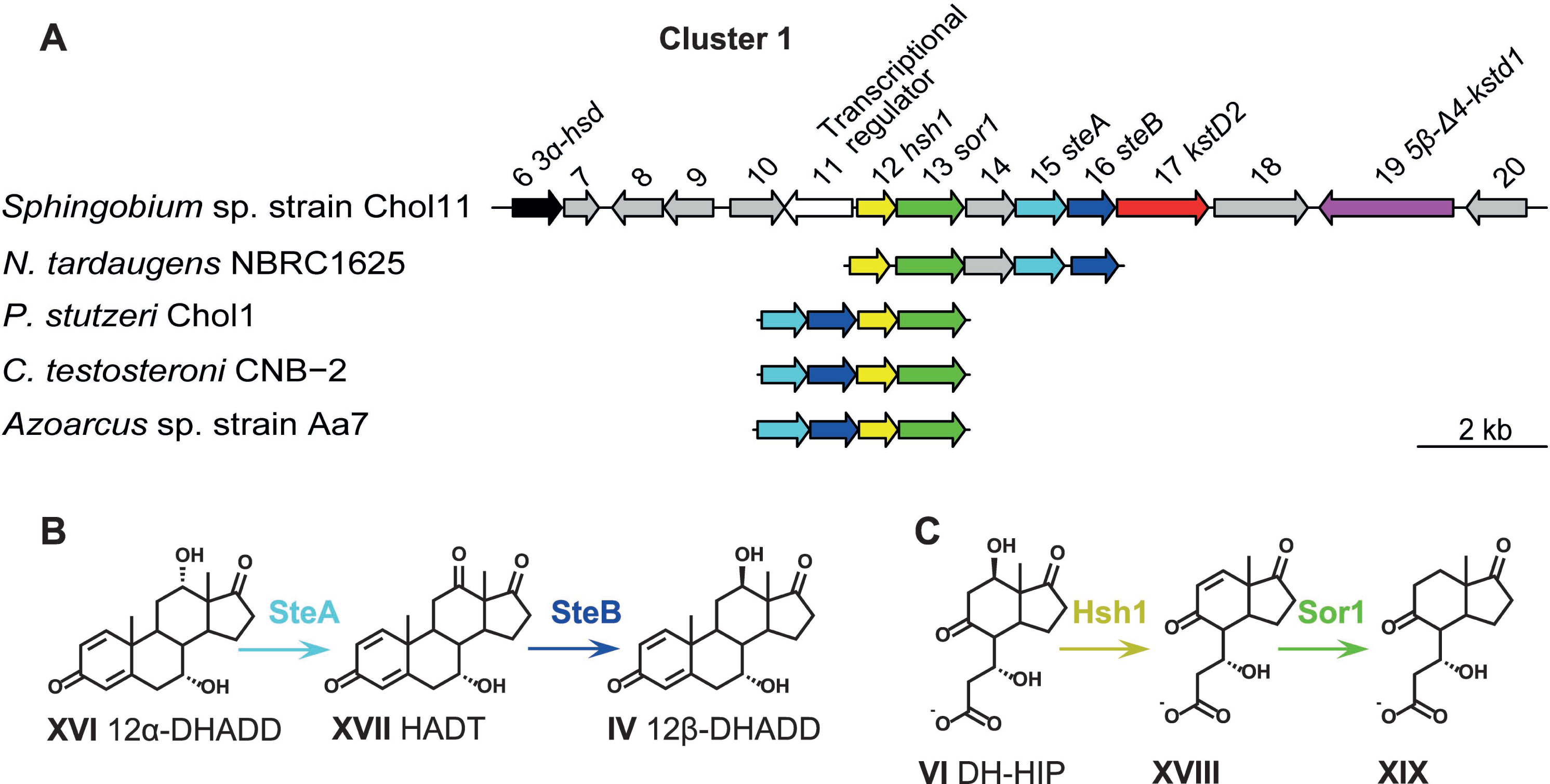
(A) Steroid degradation gene cluster 1 (*nov2c06–20*) of *Sphingobium* sp. strain Chol11 as well as similar clusters from other bile-salt degrading strains. (B) Stereoinversion of the 12-OH of the cholate-degradation intermediate DHADD as found in *C. testosteroni* (38). (C) Removal of the 7-OH of cholate-degradation intermediate DH-HIP, which corresponds to the 12-OH in cholate (23). Compound abbreviations: XVI, 12α-DHADD (7,12α-dihydroxy-androsta-1,4-diene-3,17-dione); XVI, HADT (7-hydroxy- adrosta-1,4-diene-3,12,17-trione); XVIII, 3-hydroxy-H-methyl-hexahydro-indanone-6- propenoate; XIX, 3-hydroxy-H-methyl-hexahydro-indanone-propanoate.

### 5β-Δ4-3-Ketosteroid dehydrogenase

The next step is the introduction of a double bond in the A-ring by 5β-Δ^4^-3-ketosteroid dehydrogenase (5β-Δ^4^-KSTD, named 5β-Δ^4^- KSTD1) (28). 5β-Δ^4^-KSTD1 is encoded in cluster 1 (Nov2c19) and is much more abundant in steroid- versus glucose-grown cells (>5.5-fold, Fig 4).

### 7α-Hydroxysteroid dehydratase

The key enzyme of the Δ^4,6^-pathway in strain Chol11, the 7α-hydroxysteroid dehydratase Hsh2 (Nov2c400), introduces a double bond in the B-ring by elimination of water (27). The *hsh2* gene is located in close proximity to cluster 3. Hsh2 was previously shown to be active in glucose-grown cells (28), in agreement with its similar abundance in all cells (Fig 4).

### Δ^1^-3-Ketosteroid dehydrogenase

In the Δ^1,4^-variant bile-salt degradation pathway, the next step is the introduction of a second double bond in the A-ring at C1 by a Δ^1^-3- ketosteroid dehydrogenase (Δ^1^-KSTD), which is a structural prerequisite for subsequent cleavage of the B-ring. The formation of Δ^1,4,6^-intermediates such as HATD (VIII in Fig 1) during growth with bile salts shows, that this reaction also occurs in strain Chol11 (10). Putative Δ^1^-KSTDs (36, 37) are Nov2c82 (encoded between gene clusters 1 and 2, not detected in any condition), and Nov2c695 (encoded close to gene cluster 4, about 2- to 3-fold increased abundance in steroid-grown cells) (Fig 4). This expression pattern is in line with the biotransformation assays showing no Δ^1^-KSTD activity in glucose-adapted cells. As no Δ^1,4,6^-intermediates with a side chain were reported for strain Chol11 during growth on bile salts (10, 27), this step might occur after side-chain degradation in strain Chol11.

### Oxidation of the 12-OH-group

The previously observed formation of the 3,12-dioxo- chol-4,6-dienoate (DOCDA, XIII in Fig 3) (10, 27) suggests the involvement of a 12α- dehydrogenase in the degradation of 12-hydroxy bile acids in strain Chol11. A similar reaction is catalyzed in *C. testosteroni* by SteA, which is active on 12α-hydroxy steroids without a side chain (38). In *C. testosteroni*, the resulting 12-oxo-steroids are then reduced to the corresponding 12β-steroids by SteB (Fig 5B) before cleavage of the A-ring. The presence of a 12β-OH in the degradation intermediate HATD (VIII in Fig 1) indicates that the reduction to a 12β-OH also takes place in strain Chol11. Homologs to SteA and SteB are encoded in cluster 1 (Nov2c15 and Nov2c16, respectively) and were detected with higher Mascot scores in steroid-grown cells (Fig 4).

### Phase 2: Side-chain degradation (green section in Fig 3) CoA-activation of the side chain

The steroid-C24-CoA-ligase SclA catalyzing the initial step of side chain degradation in strain Chol11 was previously described (29) and is encoded in the putative side-chain degradation gene cluster 2. SclA abundance, relative to glucose-grown cells, was 3.3-fold higher in cholate- and deoxycholate-grown cells, and 1.3-fold higher in 12β-DHADD-grown cells (Fig 4).

### Desaturation of the side chain

Previous enzyme assays with cell free extract of strain Chol11 indicated that a double bond is introduced into the CoA activated side chain (29). This reaction is typically catalyzed by α_2_β_2_-heterotetrameric acyl-CoA dehydrogenases (ACADs) during the degradation of steroids (39). Two predicted ACAD proteins in gene cluster 2 (Nov2c221 and Nov2c222) had 4- to 5-fold increased abundances in cholate- and deoxycholate-grown cells, but not in 12β-DHADD-grown cells (Fig 4). This suggests that they comprise an ACAD involved in side-chain degradation of bile salts, but the location of any double bond formed by this ACAD is unknown.

### Further side-chain degradation

The next step in bile-salt side-chain degradation is typically the addition of water to the double bond (Fig 6B). The enzymes catalyzing this hydration belong to the MaoC family of the thioesterase superfamily, containing a hotdog fold domain, or the Crotonase family (19, 40–42). These hydratases consist either of a single protein, such as the C_5_ side-chain hydratases of *P. stutzeri* Chol1 and *tuberculosis* (19, 42), or of two subunits, such as the C_3_ side-chain hydratase of *M. tuberculosis* (40). Two proteins from the thiolase superfamily with hotdog fold domains are encoded in cluster 2 (Nov2c219 and Nov2c220) adjacent to the ACAD genes. Nov2c219 was detected in only cholate- and deoxycholate-grown cells, and Nov2c220, only in cholate-grown cells (Fig 4). However, these proteins show only very low similarity (less than 20 % identity) to known side-chain hydratases, which suggests a different function for these proteins.

**Fig 6.**
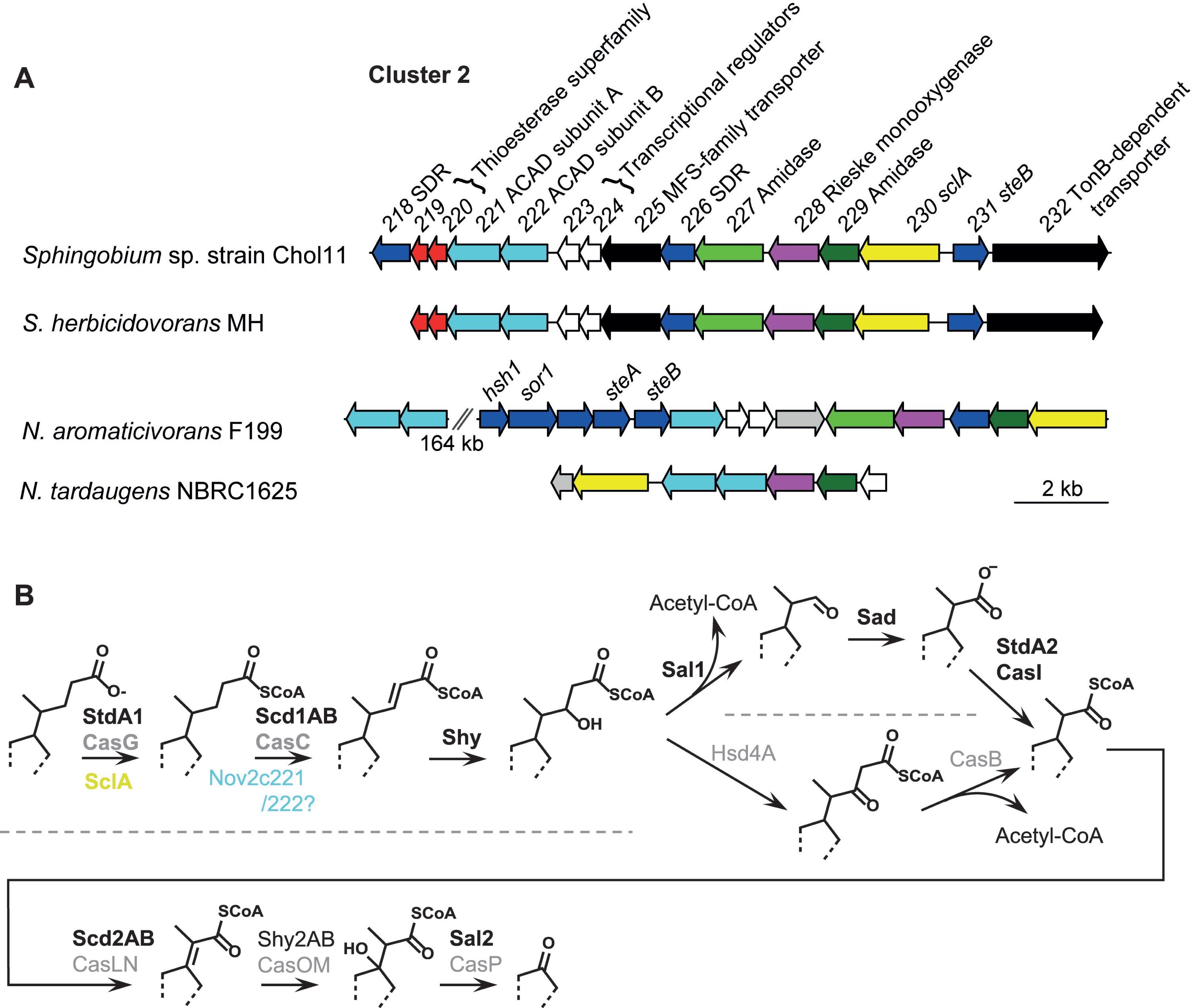
(A) Steroid degradation gene cluster 2 (*nov2c218–232*) of *Sphingobium* sp. strain Chol11 and similar clusters from other bile-salt degrading strains. (B) Bile-salt side chain degradation mechanism in *P. stutzeri* Chol1 (black enzyme names) (19, 21, 23, 24) and *R. jostii* RHA1 (grey enzyme names) (13). Colored: Corresponding enzymes in strain Chol11 (colors correspond to genes in A), bold: experimentally verified, light: suggested by bioinformatic analyses.

Homologs to other known side-chain degradation enzymes, such as thiolases and aldolases, are not encoded in cluster 2 (29). In *P. stutzeri* Chol1 and *R. jostii* RHA1, the first cycle of side-chain degradation leads to the release of acetyl-CoA and a shortened C_3_-side chain. For the degradation of the C_3_-side chain, a second cycle of aldolytic cleavage with similar steps including introduction of a double bond, addition of water and aldolytic cleavage is necessary. For this second cycle, both organisms encode a second set of enzymes (13, 19, 23).

Our proteome and bioinformatic analyses of strain Chol11 revealed no enzymes that could potentially be involved in side-chain cleavage or degradation of the C_3_-side chain. This suggests that side-chain degradation in strain Chol11 is a mechanism other than aldolytic or thiolytic cleavage. A so-far unknown alternative mechanism might involve other proteins encoded in cluster 2, including putative hydroxysteroid dehydrogenases, amidases, and a Rieske monooxygenase (Fig. 6).

### Phase 3: B-ring cleavage by the monooxygenase KshAB (yellow section in Fig 3)

The first step in the degradation of the steroid nucleus is the cleavage of the B-ring by the KshAB monooxygenase system (9, 43). Five homologs of the oxygenase component, KshA, are encoded on chromosome 2 (Nov2c66, Nov2c228, Nov2c407, Nov2c430, and Nov2c440 with 27 % - 32 % identity to KshA of *P. stutzeri* Chol1) (Fig 4). The numerous B-ring cleaving KshA homologs in the genome of strain Chol11 strongly suggest that steroid nucleus degradation starts with 9,10-*seco* cleavage, although the resulting 9,10*-seco* steroids have so far never been detected in cell-free supernatants of strain Chol11 cultures. Similar multiplicity of KshA homologs is also known from steroid degrading *Rhodococci* (44, 45). Nov2c228, Nov2c407, Nov2c430, and Nov2c440 had increased abundances in steroid-grown cells and therefore are candidates for this reaction. However, Nov2c228 has the lowest similarity to KshA_Chol1_, its encoding gene is localized in the side-chain degradation gene cluster, and the abundance of Nov2c228 was only increased in bile-salt grown cells, but not in 12β-DHADD-grown cells. Thus, a different role for this enzyme appears feasible. In this context, the similarities of the KshA oxygenases and Neverland oxygenases, that are involved in the production of ecdysteroids in arthropods (46), could indicate a wider function for Rieske monooxygenases in steroid metabolism.

Interestingly, there are no distinct homologs of the reductase component KshB encoded in the genome of strain Chol11. This was also reported for *N. tardaugens* NBRC16725 (47). A flavodoxin reductase Novbp123 with 29 % identity to KshB of *P. stutzeri* Chol1 is encoded on plasmid pSb of strain Chol11 (30). This enzyme was detected in the membrane protein-enriched fractions of all tested cells with similar Mascot scores, indicating that its synthesis is not regulated in response to steroid degradation.

Novbp123 is encoded in a gene cluster together with a ferredoxin, a cytochrome c, and several exported and membrane proteins, suggesting that it is involved in membrane electron transport rather than bile-salt degradation specifically (Fig S3). This points at a different, KshB-independent electron shuttling mechanism for the KshA homologs in strain Chol11.

### Phase 4: Complete degradation of the 9,10-*seco* intermediates (orange section in Fig. 3)

Most proteins required for the degradation of the 9,10-*seco* degradation intermediates, derived from both cholate and deoxycholate, and the respective HIP intermediates (26) are encoded in gene cluster 3 (Fig 7A). All of these proteins are at least 1.5-fold increased in abundance in steroid-grown cells (Fig. 4). This confirms that degradation of the steroid nucleus proceeds via the 9,10-*seco* pathway. Cluster 3 is very similar to the cluster encoding testosterone degradation in *N. tardaugens* NBRC16725 (47). In both organisms, genes encoding homologs for the reductase component TesA1 of the 9,10- *seco* steroid monooxygenase, TesA1A2, are missing, which could be a further hint at a different electron shuttling mechanism. However, in cluster 3 of strain Chol11, a flavin reductase (Nov2c347) is encoded near the gene for the oxygenase component, TesA2 (Nov2c349), indicating Nov2c347 could serve as a TesA1 substitute. Regarding HIP- degradation, a homolog for the gene encoding HIP-CoA-ligase ScdA (48) is missing in strain Chol11, but the CoA transferase, Nov2c359, could have this function.

**Fig 7.**
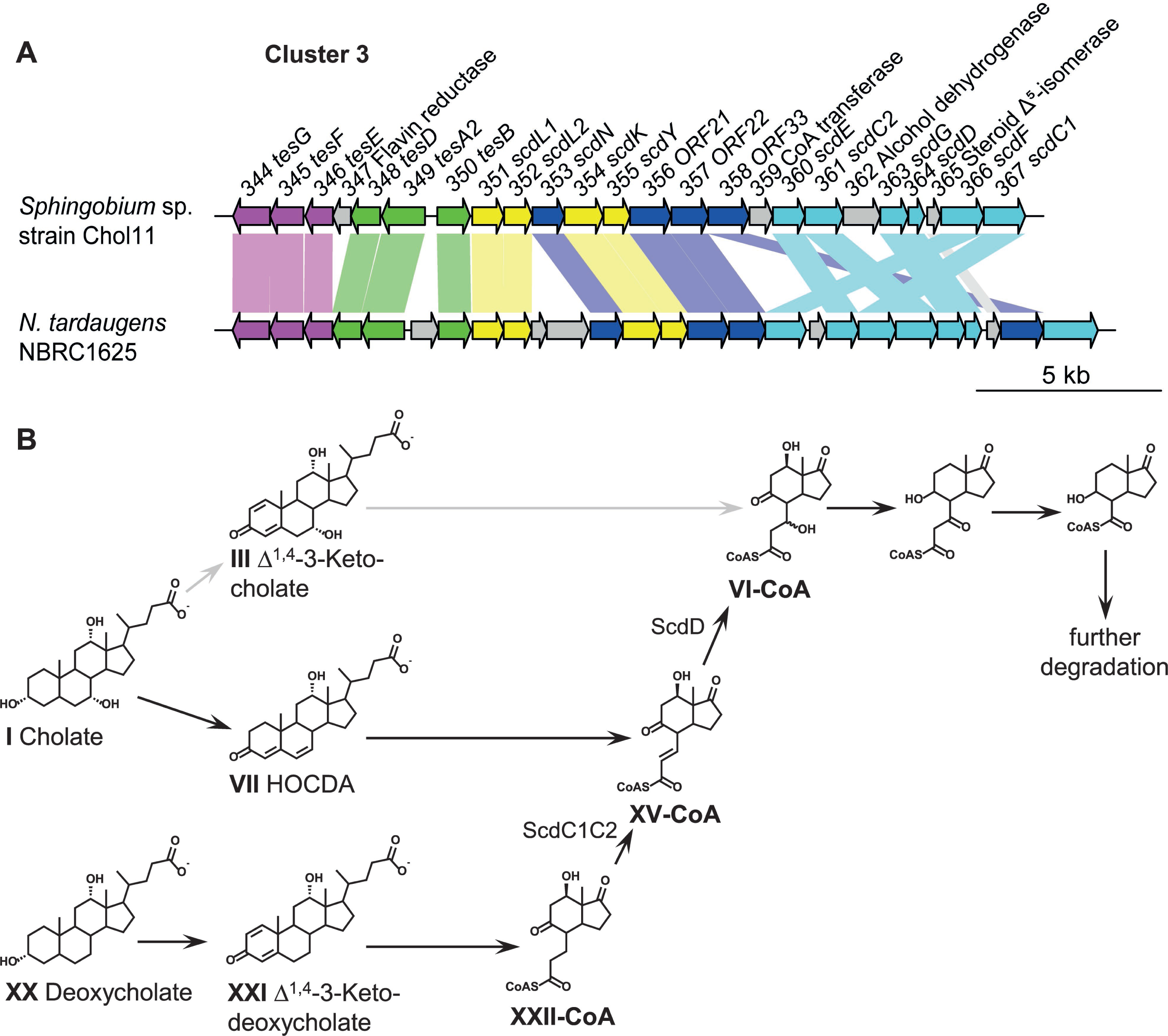
(A) Steroid degradation gene cluster 3 (*nov2c344–370*) of *Sphingobium* sp. strain Chol11 compared to the testosterone degradation gene cluster of *N. tardaugens* NBC16725 (47). Connection lines indicate homologs. (B) Proposed degradation of 7- hydroxy and 7-deoxy bile-salts in strain Chol11 (black) as opposed to degradation of 7- hydroxy bile salts via the Δ^1,4^-variant (grey). Compound abbreviations: XXII, 7-hydroxy- H-methyl-hexahydro-indanone-propanoate.

### Fate of the 12-OH

In *P. stutzeri* Chol1, the 12β-OH is removed during C- and D-ring degradation by the elimination of water catalyzed by Hsh1 and subsequent reduction of the resulting double bond by Sor1 (23) (Fig 5C). In strain Chol11, homologs to Hsh1 and Sor1 are encoded in cluster 1 near *steA* and *steB* (*nov2c12* and *nov2c13*, respectively, Fig. 5A) and were found in increased abundances in steroid-grown cells (Fig 4). A gene cluster with the same order of genes for the 12-OH transforming enzymes SteA, SteB, Hsh1 and Sor1 is present in *N. tardaugens* NBRC16725 and a similar cluster is present in *C. testosteroni* CNB-2*, P. stutzeri* Chol1, and *Azoarcus* sp. strain Aa7 (Fig 5A), implicating a general role of these enzymes in bile-salt degradation.

### Channeling of 7-hydroxy and 7-deoxy bile salts into C- and D- ring degradation

During degradation of HIP-intermediates (such as VI, XV, and XXII in Fig 7), the remainder of the B-ring is degraded via β-oxidation, which requires a hydroxy group at the former C7 (23) (Fig 7B). This hydroxy group is present in 7-hydroxy bile salts such as cholate, but during the degradation of 7-deoxy bile salts, such as deoxycholate, it has to be introduced into the propanoate side chain of the respective HIP-intermediates. This is initiated by the introduction of a double bond by the heteromeric ACAD ScdC1C2 followed by the addition of water by the hydratase ScdD (23, 49, 50).

In Δ^4,6^-intermediates, the hydroxy group at C7 is eliminated. Thus, bile salt degradation by the Δ^4,6^-pathway variant results in HIP-like intermediates with a double bond in the propanoate side chain attached to ring C (XV). This is the same intermediate as found during degradation of 7-deoxy bile salts after introduction of a double bond (23), and the needed hydroxy group could be added by the hydratase, Nov2c364, which was detected all steroid grown cells in similar abundance (Fig 4). Although the ACAD reaction catalyzed by ScdC1C2 is not needed for the degradation of cholate via the Δ^4,6^-variant, both subunits, Nov2c367 and Nov2c361, were found in all steroid-grown cells with similar Mascot scores (Fig 4). Thus, it is possible that the Δ^6^-double bond had meanwhile been reduced, necessitating the ACAD.

As the hydroxy group eliminated by Hsh2 must again be added at the stage of HIP intermediates, the benefit of the elimination remains unclear. It might be related to the fact that many intermediates of bile-salt degradation are excreted in significant amounts during growth of bile-salt degrading bacteria not only in laboratory cultures but also in soil samples (10, 34). Other bile-salt degrading strains, such as *P. stutzeri* Chol1, that exclusively use the Δ^1,4^-degradation pathway are unable to utilize Δ^4,6^-compounds as growth substrates (10, 27). To this end, the dehydration might provide a way for strain Chol11 to secure the individual availability to these important carbon sources in natural habitats.

### The Δ^4,6^-variant is widespread within the family *Sphingomonadaceae* Prediction of the Δ^4,6^-variant pathway in steroid degrading bacteria

Apart from strain Chol11, steroid degradation has been reported in other sphingomonads such as *tardaugens* NBRC16725 (47, 51) and *Sphingomonas* sp. strain KC8 (52). In addition, *Sphingomonas* sp. strain Chol10 was also isolated as a HOCDA-degrading bacterium (10).

To investigate the prevalence of the Δ^4,6^-variant pathway within the family *Sphingomonadaceae*, we searched all 398 complete and draft genomes from the genera *Sphingobium, Novosphingobium,* and *Sphingomonas* available from the NCBI RefSeq database for the simultaneous presence of key steroid degradation proteins and Hsh2, the key protein of the Δ^4,6^-pathway. First, steroid degrading bacteria were predicted using 23 Hidden Markov Models (HMMs) representing ten key proteins of canonical steroid nucleus degradation (53). Based on the presence of seven out of ten key proteins including KshA and TesB, 53 genomes were predicted to encode steroid degradation. Second, Hsh2 orthologs were determined in these genomes by BLASTp analyses using Hsh2 of *Sphingobium* sp. strain Chol11 (27) as query. Thirty-nine genomes containing both, the steroid-degradation genes and *hsh2,* were found. To further confirm the prediction of these proteins being involved in steroid degradation in these organisms, a reciprocal BLASTp analysis was performed using steroid degradation proteins from *P. stutzeri* Chol1 and Hsh2_Chol11_ as queries (Fig 8). Thirty- eight genomes were confirmed to encode Hsh2 orthologs, and all of them contained orthologs for the majority of key proteins for steroid nucleus degradation. However, only a small subset of side-chain degradation proteins is encoded in most of the *Sphingobium* and *Novosphingobium* genomes. This was in contrast to the *Sphingomonas* genomes where orthologs of the genes encoding CoA-ligase, heteromeric ACAD, heteromeric hydratase, and aldolase for the degradation of the C_3_- side chain from *P. stutzeri* Chol1 are present.

**Fig 8.**
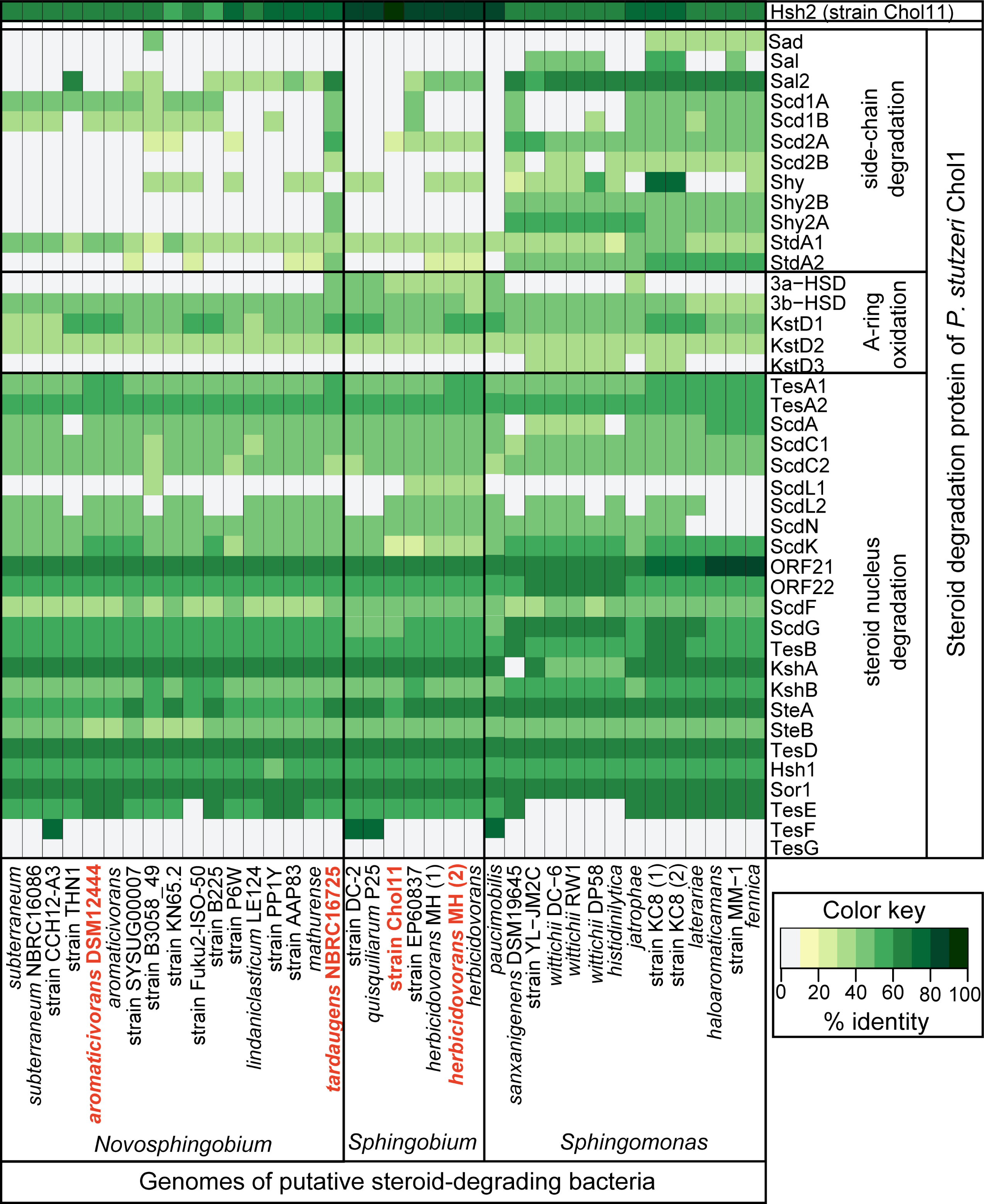
Distribution of steroid degradation gene orthologs in 38 *Sphingomonas, Sphingobium,* and *Novosphingobium* strains. Heat-map showing the BLAST identities for reciprocal BLASTp hits to Hsh2 from *Sphingobium* sp. strain Chol11 and the bile-salt degradation proteins from *P. stutzeri* Chol1. Names of strains tested and found to use the Δ^4,6^-variant pathway are in red. For accession numbers of the genomes see Fig S4.

This suggests that bile-salt degradation via Δ^4,6^-intermediates is widely distributed among members of *Sphingomonas, Sphingobium,* and *Novosphingobium*, while the distinct side-chain degradation mechanism proposed for strain Chol11 may occur in most members of *Sphingobium* and *Novosphingobium*.

### Bile-salt degradation in strains predicted to use the Δ^4,6^-variant

To investigate whether strains predicted to degrade bile salts via the Δ^4,6^-pathway variant, a selection of type strains with complete genome sequences was analysed for bile salt degradation. *Novosphingobium aromaticivorans* F199, *Sphingobium herbicidovorans* MH, and *N. tardaugens* NBRC16725 were tested for the degradation of the 7α-hydroxy bile salts, cholate and chenodeoxycholate, and the 7-deoxy bile salt, deoxycholate (structures in Fig 9). Accumulation of Δ^4,6^-intermediates was also monitored.

**Fig 9.**
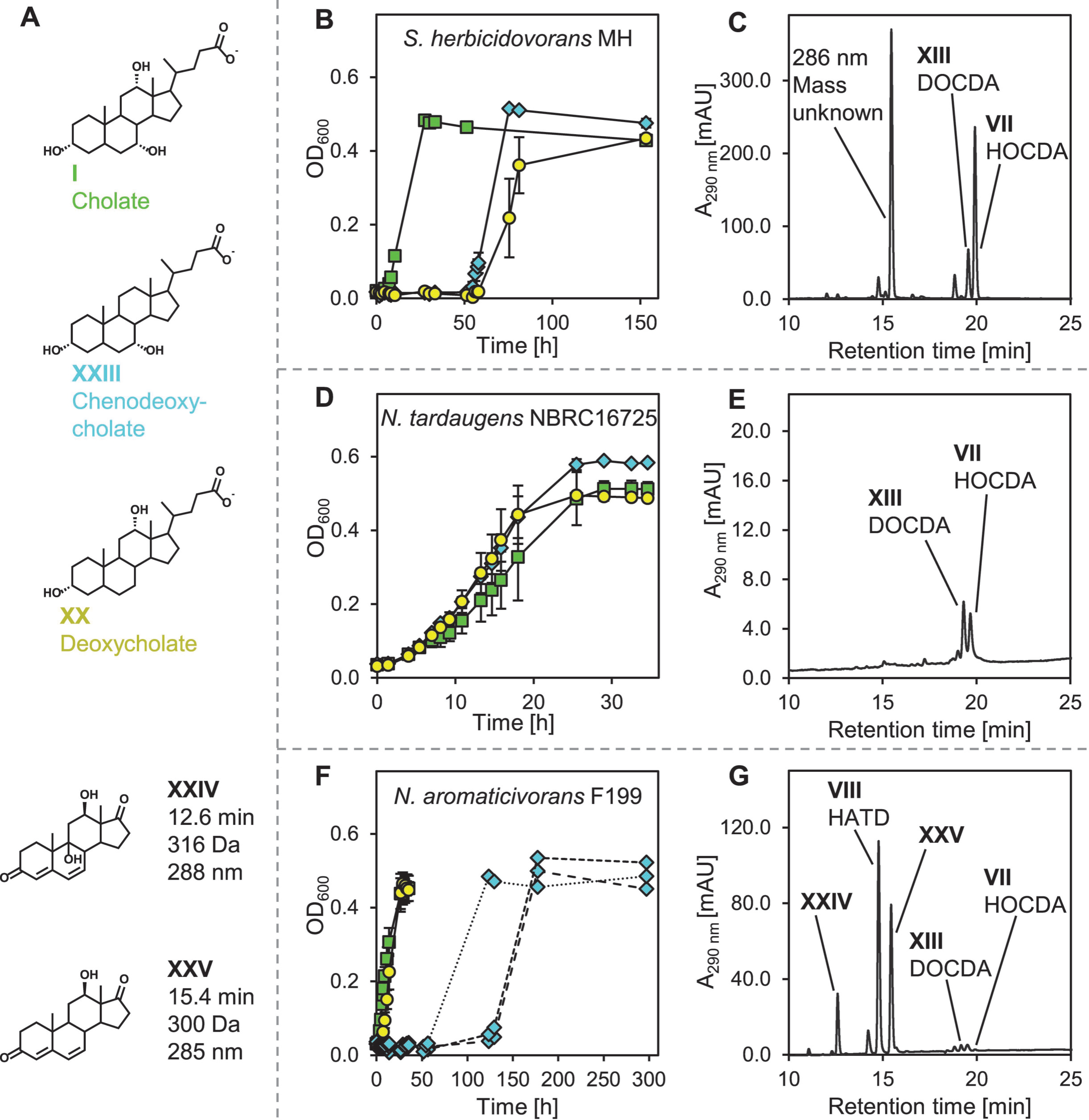
Degradation of bile salts by selected strains predicted to use the Δ^4,6^-variant of the steroid degradation pathway. (A) Structures of the tested bile salts and predicted intermediates detected in cell-free culture supernatants. (B,D,F) Growth on 1 mM bile salts. Colors of symbols correspond to colors of substrate names. Values are means of triplicates with standard deviation error bars, except in F where replicates on chenodeoxycholate are plotted separately. (C,E,G) Accumulation of intermediates during degradation of cholate in culture supernatants of the respective strains. See Fig 3 for structures not shown here. HPLC-UV chromatograms at 290 nm of culture supernatants. Steroid compounds were identified by retention time, absorbance spectrum and mass. Compound abbreviations: XXIV, 9,12-dihydroxy-androsta-4,6-diene-3,17-dione; XXV: 12-hydroxy-androsta-4,6-diene-3,17-dione.

Strains MH, NBRC16725 and F199 grew on all tested bile salts (Fig 9B,D,F), and transiently accumulated the characteristic Δ^4,6^-intermediates HOCDA (VII in Fig 1) and DOCDA (XIII in Fig 3) (Fig 9C,E,G,I). In addition, during degradation of the 7α-hydroxy bile salts, strain F199 formed the intermediates XXIV and XXV that also have a Δ^4,6^- structure and lack a side chain according to their UV- and mass spectra, respectively (Fig 9I). This supports our prediction that the Δ^4,6^-variant is the predominant pathway for bile acid degradation in *Sphingobium,* and *Novosphingobium* strains.

Like strain Chol11, strains NBRC16725, MH, and F119 were able to fully metabolize side-chain bearing bile salts, despite encoding not all sidechain catabolic enzymes proteins known from other bile salt degrading bacteria. A BLASTp analysis revealed the presence of gene clusters with high similarity to gene cluster 2 of strain Chol11 in all three strains (Fig 6A); although the clusters from the *Novosphingobium* strains, NBRC16725 and F119, were notably missing several genes. In particular, homologs of the thiolase superfamily proteins Nov2c219 and Nov2c220, the only candidates for side chain hydratases in strain Chol11, were absent in the genome of NBRC16725. Homologs of SclA, the putative ACAD Nov2c221/Nov2c222, the Rieske monooxygenase Nov2c228, and the putative amidases Nov2c227 and Nov2c229 were found in the predicted side-chain degradation clusters of all three strains.

### Degradation of conjugated bile salts

Strain Chol11 grew on the conjugated bile salts, taurocholate and glycocholate (X and XI, respectively, in Fig. 3, but not on the free amino acids glycine or taurine (Fig 10B,E). The inability to grow on these amino acids agrees with the similar biomass yields on taurocholate and glycocholate versus on unconjugated cholate (27). These results suggests that glycine or taurine are removed by amidases prior to metabolism of the free bile salts. Two candidate amidases are Nov2c227 and Nov2c229), encoded in the predicted side-chain degradation gene cluster, were 2.3- to 8.9-fold more abundant in cholate- and deoxycholate-grown cells.

**Fig 10.**
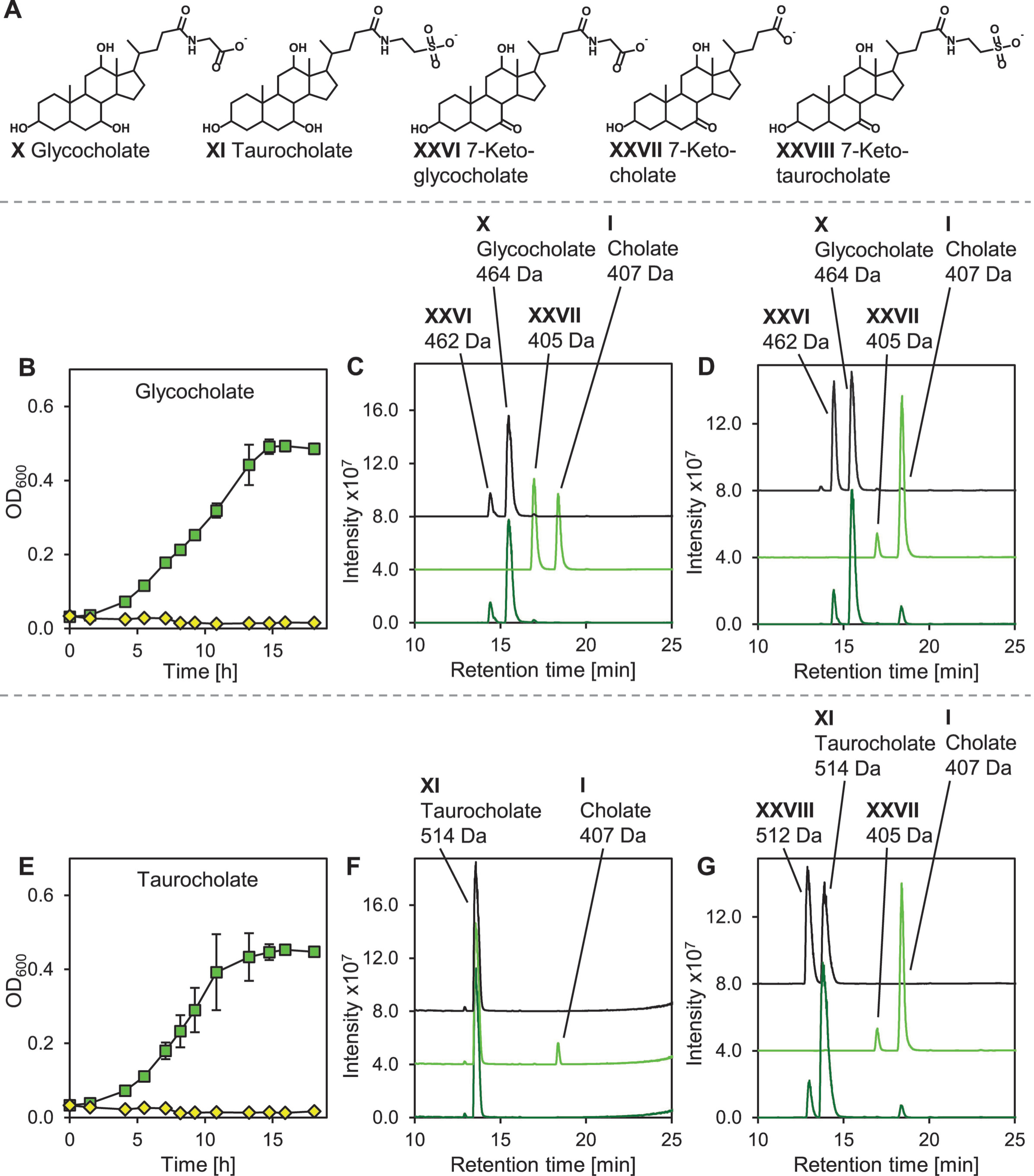
Utilization of conjugated bile salts by *Sphingobium* sp. strain Chol11 and transformation of conjugated bile salts by the amidases Nov2c227 and Nov2c229. (A) Structures of conjugated bile salts and 7-keto metabolites. (B) Growth of strain Chol11 on 1 mM glycocholate (green squares) or glycine (yellow diamonds). (C) Transformation of glycocholate by whole cells for 11 days, and (D) by cell-free extracts for 15 h. In C, D, F and G: black trace, *E. coli* MG1655 pBBR1MCS-5 (empty vector control); light green,*E. coli* MG1655 pBBR1MCS-5::*nov2c227*; and dark green, *E. coli* MG1655 pBBR1MCS- 5::*nov2c229*. (E) Growth of strain Chol11 on 1 mM taurocholate (green squares) and taurine (yellow diamonds). (F) Transformation of taurocholate by whole cells for 4 days and (G) by cell-free extracts for 15 h. Error bars indicate standard deviation (n=3). HPLC-MS base peak chromatograms measured in negative ion mode. Steroid compounds were identified by retention time, absorbance spectrum and mass. Masses are indicated for the respective deprotonated acids.

To test their potential role in deconjugation of bile salts, Nov2c227 and Nov2c229, were each heterologously expressed in *E. coli* MG1655. Cell suspensions and cell-free extracts of *E. coli* expressing Nov2c227 transformed both taurocholate and glycocholate to cholate (Fig 10). Cell suspensions and cell-free extracts of *E. coli* expressing the other amidase candidate, Nov2c229, as well as the empty vector control did not substantially catalyze this transformation. Additionally, compounds 2 Da lighter than the conjugated bile salts and cholate were present in almost all assays. *E. coli* possesses a 7α-hydroxy steroid dehydrogenase, which catalyzes the oxidation of the 7α-OH of conjugated and free bile salts (54). Therefore, it is likely that the additional compounds are the 7-keto- derivatives of the conjugated bile salts and cholate (XXVI, XXVII, and XXVIII in Fig 10). Since Nov2c227 cleaves the conjugated bile salts, glycocholate and taurocholate, it was named **B**ile **S**alt **A**midase (Bsa).

Interestingly, Bsa contains a TAT signal peptide as predicted by SignalP, indicating that this protein is secreted, and deconjugation of conjugated bile salts takes place in the periplasmatic or extracellular space prior to bile-salt uptake. In contrast to the well- elucidated N-terminal nucleophile family bile-salt hydrolases from probiotic lactic acid bacteria such as *Bifidobacterium longum* (55), Nov2c227 belongs to the large family of amidases. This indicates a different evolutionary origin for bile-salt hydrolases in steroid degradation as opposed to those in intestinal bacteria, despite their common function. Homologs of Bsa are encoded in genomes of many strains of the family *Sphingomonadaceae*, e.g. EGO055_026080 from *N. tardaugens* NBRC16725. Furthermore, *C. testosteroni* KF-1 was reported to degrade taurocholate (56) and its genome encodes two Bsa homologs (identity for both 44 %). The respective homologs, ORF25 and ORF26, from model organism *C. testosteroni* TA441 are encoded in its steroid degradation mega-cluster (14, 26). The function of Nov2c229 remains unknown so far.

## Conclusion and general discussion

Differential proteome analyses of *Sphingobium* sp. strain Chol11 together with bioinformatic analyses revealed a comprehensive set of candidate proteins for the complete degradation of bile salts. From these data the complete degradation pathway for the steroid nucleus could be deduced, which is a mosaic of unknown reactions (especially side-chain degradation) and reactions known from other steroid-degrading organisms (Fig 3). Apparently, despite the variation in the first steps of 7-hydroxy bile salt degradation in strain Chol11, which leads to the introduction of a double bond in the B-ring by water elimination catalyzed by Hsh2, further degradation of the Δ^4,6^-intermediates is very similar to the 9,10-*seco* pathway known from other organisms and steroids. However, the Δ^4,6^-variation produces steroid intermediates that cannot be utilized by other organisms such as *P. stutzeri* Chol1 (10), suggesting the involvement of specialized enzymes in organisms that are able to utilize Δ^4,6^-intermediates. The prediction and identification of further organisms degrading bile salts via this Δ^4,6^-variant of the 9,10-*seco*-pathway demonstrates a wide distribution of this variant in the family *Sphingomonadaceae*.

In addition, the interpretation of the proteome analyses in conjunction with bioinformatic analyses resulted in the identification of a side-chain degradation gene cluster encoding key proteins that are presumably responsible for side chain removal by a yet unknown mechanism. The bile-salt hydrolase activity of Bsa as well as the CoA-ester formation by the CoA ligase SclA (29) encoded in this gene cluster further corroborates the involvement of this cluster in bile salt side-chain degradation. The subsequent introduction of a double bond into a yet unknown position is presumably catalyzed by the heteromeric ACAD Nov2c221/222. The absence of several other side-chain degradation genes from the genomes of strain Chol11, *N. tardaugens* NBRC16725, and *N. aromaticivorans* F199 suggests a mechanism different from thiolytic and aldolytic cleavage. Interestingly, homologs of SclA, the putative ACAD Nov2c221/Nov2c222, the Rieske monooxygenase Nov2c228, and the putative amidase Nov2c229 were found in all Sphingomonads confirmed to use the Δ^4,6^-variant for bile salt degradation. However, the functions of this Rieske monooxygenase and the putative amidase during bile salt side-chain degradation remain unclear. In addition to being highly conserved and encoded near to confirmed side-chain degradation genes in all four tested strains, they seem to be formed exclusively in response to side-chain containing bile salts in strain Chol11. Further investigations regarding this gene cluster using molecular methods are under way. Unfortunately, these analyses are impaired by the complicated genetic modification of strain Chol11.

Interestingly, several members of the family *Sphingomonadacea* are adapted to growth with steroids and bile salts. Moreover, both *Sphingobium* sp. strain Chol11 and *N. tardaugens* NBRC16725 grow only slowly with non-steroidal substrates (51) and some genes for early steps of bile-salt degradation are apparently constitutively induced.

Together with the prevalence of bile-salt degradation in strains that had originally been isolated with xenobiotic compounds (57, 58) indicates that bile-salt degradation may be a conserved property of these organisms and calls attention to its evolutionary origin.

### Experimental procedures Cultivation of bacteria

If not indicated otherwise, *Sphingobium* sp. strain Chol11 (DSM 110934) (10), *S. herbicidovorans* MH (DSM 11019) (58), *N. aromaticivorans* F199 (DSM12444) (59), *N. tardaugens* NBRC16725 (DSM 16702) (51), and *E. coli* MG1655 (DSM 18039) (60) were cultivated in HEPES-buffered medium B (MB) (61). If not indicated otherwise, wildtype strains other than *E. coli* were grown with 1 mM cholate as the sole carbon source, whereas *E. coli* MG1655 was grown with 15 mM glucose. For maintenance, strain Chol11 and *N. tardaugens* NBRC16725 were grown on MB agar with 1 mM cholate, *S. herbicidovorans* MH was grown on CASO agar (Merck Millipore, Burlington, MA, USA), *N. aromaticivorans* F199 was grown on MB agar with 15 mM glucose, and *E. coli* MG1655 was grown on LB agar (62). For cultivation of strains containing pBBR1MCS-5 (63), 20 μg ml^−1^ gentamicin was added. When bile salts were added, gentamicin was omitted. Liquid cultures with volumes up to 5 ml were cultivated in 10 ml test tubes at 200 rpm, larger cultures were cultivated in 500 ml Erlenmeyer flasks without baffles at 120 rpm. All strains were incubated at 30 °C, except for strain maintenance of *E. coli* strains (37 °C). For agar plates, 1.5 % (w/v) Bacto agar (BD, Sparks, USA) was added.

Cholate (≥99 %), deoxycholate (≥97 %), and glycocholate (≥97 %) were purchased from Sigma-Aldrich (St. Louis, MO, USA). Chenodeoxycholate (≥98 %) was purchased from Carl Roth (Karlsruhe, Germany). Taurocholate (>98 %) was purchased from Fluka (Buchs, Switzerland).

### Growth experiments

For growth experiments, 3 – 5 ml main cultures were inoculated to a predefined OD_600_ (about 0.02) directly from liquid starter cultures and growth was determined by measurement of OD_600_ (Camspec M107, Spectronic Camspec, Leeds, UK). Growth with bile salts was tested using 1 mM cholate, 1 mM chenodeoxycholate or 1 mM deoxycholate. Growth with conjugated bile salts was tested using 1 mM taurocholate, 1 mM glycocholate, 15 mM glycine or 15 mM taurine. At suitable time points, samples for HPLC-MS measurements were withdrawn.

Starter cultures were grown with 1 mM cholate for strain Chol11, *N. aromaticivorans* F199, and *N. tardaugens* NBRC16725 or 12 mM succinate for *S. herbicidovorans* MH. Starter cultures were incubated over-night for about 15 h.

### Biotransformation experiments

Induction of cholate degradation in strain Chol11 was tested using suspensions of cholate- and glucose-grown cells with 1 mM cholate and 10 μg ml^-1^ chloramphenicol to inhibit *de novo* protein synthesis as described (28).

For determining whole-cell biotransformation of various steroid compounds by *E. coli* MG1655 expressing amidase genes, 5 ml starter cultures of *E. coli* MG1655 pBBR1MCS-5 as empty vector control, *E. coli* MG1655 pBBR1MCS-5::*nov2c227* or *E. coli* MG1655 pBBR1MCS-5::*nov2c229* in LB with 20 μg ml^-1^ gentamicin were incubated for about 15 h. Cells were harvested by centrifugation in 2-ml reaction tubes (8,000 x g, ambient temperature, 3 min), washed with MB and resuspended in MB to an OD_600_ of about 1. Cell suspensions were incubated for several days at 30 °C at 200 rpm after addition of either 1 mM taurocholate or 1 mM glycocholate. 30 mM glucose were added to all preparations as carbon source.

For monitoring biotransformations of conjugated bile salts by cell free extracts of *E. coli* MG1655 expressing amidase genes, 50 ml LB with 20 μg ml^−1^ gentamicin were inoculated to an initial OD_600_ of 0.015 with the aforementioned strains of *E. coli* MG1655 and incubated for about 18 h with addition of 0.2 mM isopropyl-β-D-thiogalactopyranosid after about 3 h. Cells were harvested (8,000 x g, 4 °C, 8 min), washed with 10 mM MOPS buffer (pH 7.8) and resuspended in about 2 ml 50 mM MOPS buffer (pH 7.8). Cells were disrupted in 15-ml conical centrifugation tubes by ultrasonication on ice (amplitude 60 %, cycle 0.5, UP200S, Hielscher Ultrasonics, Teltow, Germany) for 8 min with a 1 min break after 4 min. Cell debris was removed by centrifugation (25,000 x g, 4 °C, 30 min) in 2-ml reaction tubes. Cell extracts were used immediately in enzyme assays or stored at -20 °C for later use. Enzyme assays (1 ml) contained 50 mM MOPS (pH 7.8), 1 mM substrate and 100 μl cell extract. Assays were incubated for 30 min – 15 h at 30 °C. Samples of all enzyme assays were subjected to HPLC-MS measurements

## Proteome analyses

### Generation of substrate adapted cells

Strain Chol11 was freshly thawed from a cryopreservation culture for each cultivation on MB agar and streaked twice onto plates with 1 mM cholate. After 1 – 4 days, the strain was transferred to new agar plates containing either 1 mM cholate, 1 mM deoxycholate, 2 mM 12β-DHADD or 15 mM glucose. Cells were further transferred twice to the same medium after incubation for 2 days with steroidal substrates or 3 – 4 days with glucose. From these plates, twelve 5 ml MB starter cultures containing the same carbon source as the respective solid media were inoculated and incubated for about 15 h for steroidal compounds or 30 h for glucose. Subsequently, 1 mM bile salt, 2 mM 12β-DHADD or 15 mM glucose were added and the starter cultures were incubated for further 1 – 1.5 h. Starter cultures with the same substrate were pooled and used for inoculation of twelve 100 ml main cultures in 500-ml Erlenmeyer flasks without baffles containing the same carbon sources at an OD_600_ of 0.015. Main cultures were incubated at 30 °C and 200 rpm and growth was monitored until the half maximal OD_600_ was reached. Cultures were harvested in two 50- ml conical centrifugation tubes by centrifugation (5,525 x g, 4 °C, 30 min) and kept on ice. Cells were washed with 25 ml 100 mM Tris buffer (pH 7.5 with HCl) containing 5 mM MgCl_2_ and cells of the same culture were pooled. After centrifugation, cells were resuspended in 625 μl of the same buffer, transferred to 2 ml reaction tubes and harvested by centrifugation (14,000 x g, 4 °C, 15 min). After weighing, pellets were snap-frozen using liquid nitrogen and stored at -70 °C.

### Profiling of soluble proteins by 2D DIGE and protein identification by MALDI-TOF-MS/MS

Soluble proteins were extracted from cells of strain Chol11 and 2D DIGE performed essentially as described previously (64). Per growth condition, four biological replicate samples were prepared and 50 µg total protein used for minimal labelling with 200 pmol of Lightning SciDye DIGE fluors (SERVA Electrophoresis GmbH, Heidelberg, Germany). Glucose-adapted cells served as reference state and were labelled with Sci5. Protein extracts of the other three (test) states were each labelled with Sci3. The internal standard contained equal amounts of all test and the reference state(s) and was labelled with Sci2. First dimension separation by isoelectric focusing (IEF) was conducted with 24 cm-long IPG strips (pH 3–11 NL; GE Healthcare) run in a Protean i12 system (Bio-Rad, Munich, Germany). The IEF program used was as follows: 50 V for 13 h, 200 V for 1 h, 1,000 V for 1 h, gradual gradient to 10,000 V within 2 h and 10,000 V until 80,000 Vhs were reached. Second dimension separation according to molecular size was done by SDS-PAGE (12.5% gels, v/v) using an EttanDalt*twelve* system (GE Healthcare).

Directly after electrophoresis, 2D DIGE gels were digitalized using a CCD camera system (Intas Advanced 2D Imager; Intas Science Imaging Instruments GmbH, Göttingen, Germany) (65). Cropped gel images were analyzed with the SameSpots^™^ software (version 5.0.5.0, TotalLab, Newcastle upon Tyne, UK) and spots with changes in abundance of ≥ **|**1.5**|**-fold and an ANOVA p-value of ≤ 1×10^-4^ were accepted as significant (66). Spots of interest were excised from at least two separate, preparative colloidal Coomassie Brilliant Blue stained gels (300 µg protein load) using the EXQuest spot cutter (Bio-Rad), and subsequently washed and tryptically digested as described recently (67).

Sample digests were spotted onto Anchorchip steel targets (Bruker Daltonik GmbH, Bremen, Germany) and analyzed with an UltrafleXtreme MALDI-TOF/TOF mass spectrometer (Bruker Daltonik GmbH) as recently described (67). Peptide mass fingerprint (PMF) searches were performed with a Mascot server (version 2.3; Matrix Science, London, UK) against the translated genome of strain Chol11 with a mass tolerance of 25 ppm. Five lift spectra were collected to confirm PMF identification and three additional spectra were acquired of unassigned peaks applying feedback by the ProteinScape platform (version 3.1; Bruker Daltonik GmbH). In case of failed PMF identification, eight lift spectra of suitable precursors were acquired. MS/MS searches were performed with a mass tolerance of 100 ppm. For both, MS and MS/MS searches, Mascot scores not meeting the 95% certainty criterion were not considered significant. A single miscleavage was allowed (enzyme trypsin) and carbamidomethyl (C) and oxidation (M) were set as fixed and variable modifications, respectively.

### Shotgun proteomic analysis

For shotgun analysis, cell pellets of three biological replicate samples per growth condition were suspended in lysis buffer, cells disrupted and the debris-free fraction reduced, alkylated and subjected to tryptic digest as previously described (67). Obtained peptides were separated by nanoLC (UltiMate 3000; ThermoFisher Scientific, Germering, Germany) using a trap-column (C18, 5 µm bead size, 2 cm length, 75 µm inner diameter; ThermoFisher Scientific) and a 25 cm analytical column (C18, 2 µm bead size, 75 µm inner diameter; ThermoFisher Scientific) applying a 360 min linear gradient (68). The nanoLC eluent was continuously analyzed by an online-coupled ion-trap mass spectrometer (amaZon speed ETD; Bruker Daltonik GmbH) using the captive spray electrospray ion source (Bruker Daltonik GmbH). The instrument was operated in positive mode with a capillary current of 1.3 kV and drygas flow of 3 l min ^-1^ at 150°C. Active precursor exclusion was set for 0.2 min. Per full scan MS, 20 MS/MS spectra of the most intense masses were acquired. Protein identification was performed with ProteinScape as described above, including a mass tolerance of 0.3 Da for MS and 0.4 Da for MS/MS searches and applying a target decoy strategy (false discovery rate < 1%).

### Analysis of the membrane protein-enriched fraction

Total membrane fractions were prepared from two biological replicates per substrate condition as described (67). The obtained protein content was determined with the RC-DC assay (Bio-Rad) and 8 µg total protein separated by SDS-PAGE gels (∼7 cm separation gel). Following staining with Coomassie Brilliant Blue (69), each sample lane was cut into 8 slices and each slice into small pieces of ∼1-2 mm³ prior to washing, reduction, alkylation and tryptic digest (67). Separation and mass determination was performed as described above, using a 100 min linear gradient. Identified proteins (performed as described above) of each slice per sample were compiled using the protein extractor of the ProteinScape platform.

### Cloning techniques

Cloning was performed according to standard procedures and as described elsewhere (27).

For expression of amidase genes in *E. coli* MG1655, genes were amplified using the respective primer pairs expfor/exprev (Tab 1) and ligated into vector pBBR1MCS-5. The respective ligation products were transferred to *E. coli* MG1655 by heat shock transformation. Presence and correct ligation of plasmids was confirmed by colony PCR using M13 primers.

### Analytical methods

Steroid compounds were analyzed by HPLC-MS. For this, samples were centrifuged (>16,000 x g, ambient temperature, 5 min) to remove cells and particles. Supernatants were stored at -20 °C and centrifuged again prior to measurement. Samples from cell suspension experiments with *E. coli* MG1655 were directly frozen at -20 °C and only centrifuged after thawing to break the cells. HPLC-MS measurements were performed with a Dionex Ultimate 3000 HPLC (ThermoFisher Scientific, Waltham, MA, USA) with an UV/visible light diode array detector and coupled to an ion trap amaZon speed mass spectrometer (Bruker Daltonik, Bremen, Germany) with an electrospray ion source. Compounds were separated over a reversed phase C_18_ column (150 x 3 mm, Eurosphere II, 100-5 C_18_; Knauer Wissenschaftliche Geräte, Berlin, Germany) at 25 °C. Samples of cell suspensions for testing the induction of steroid degradation in *Sphingobium* sp. strain Chol11 were measured as described by (27), whereas all other samples were measured as described by (28).

Bile salt concentrations were determined as peak area from base peak chromatograms measured in negative ion mode. Intermediates were identified according to retention time, UV- and MS-spectra, and comparison with known compounds. Structures of unknown metabolites were proposed on the basis of retention time as well as UV- and MS-spectra.

### Bioinformatic methods

Searches for homologous proteins and determinations of protein similarities were performed using the BLASTp algorithm (70, 71). Protein similarities were calculated from global alignments in the BLAST suite using the Needleman-Wunsch algorithm (70, 72) Protein domains and families were predicted using Interpro (73). For functional annotation of strain Chol11, the eggNOG database (74) was used.

For the bioinformatic identification of other sphingomonads using the Δ^4,6^-variant of the9,10-*seco*-pathway, first a database of putative steroid degraders was set up similar to (75). On 18 October 2018, all complete and draft genomes available for the genera *Sphingobium, Novosphingobium,* and *Sphingomonas* were downloaded from the RefSeq database (version 89). Using 23 Hidden Markov Models (HMMs) (53), these genomes were searched for ten homologs of steroid degradation proteins using HMMER v3 (76) using a maximum *E-*value of 10^-25^. HMMs for Δ^1^-KSTD (KstD), KshA, TesA2 (=HsaA in *R. jostii* RHA1), TesB (=HsaC), TesE (=HsaE), TesF (=HsaG), TesG (=HsaF), ScdK (=IpdC), ScdL1 (=IpdA), and ScdL2 (=IdpB) were used. Bacteria were predicted to be able to degrade steroids when their genomes encoded homologs of seven out of the eleven steroid degradation key enzymes including KshA and TesA2. With these genomes, a reciprocal BLASTp search (75) was conducted using the key enzyme of the Δ^4,6^-pathway variant Hsh2 from strain Chol11 (27) as query using *E*-value and identity cutoffs of 10^-25^ and 35 %, respectively. These values were optimized empirically comparing analyses using Hsh2_Chol11_ as well as BaiE from *C. scindens* (UniProt-ID P19412) which has a similar function in a different pathway (77). The results of both analyses were compared and *E-*value and identity cutoffs were chosen to ensure, that proteins were only identified as homologs of one of these dehydratases. All genomes from putative steroid degraders containing Hsh2 homologs were subjected to a reciprocal BLASTp analysis using known steroid degradation proteins from *P. stutzeri* Chol1 as queries. For data analysis and preparation of figures, R (v3.5.1) was used together with the packages circlize (v0.4.8), genoPlotR (v0.8.9), ggplot2 (v3.2.1), ComplexHeatmap (v1.18.1), gplots (v3.0.3), RColorBrewer (v1.1-2), VennDiagram (v1.6.20), ape (v5.3), reshape2 (v1.4.3), tidyverse (v1.3.0), and readxl (v1.3.1).

**Tab 1.**
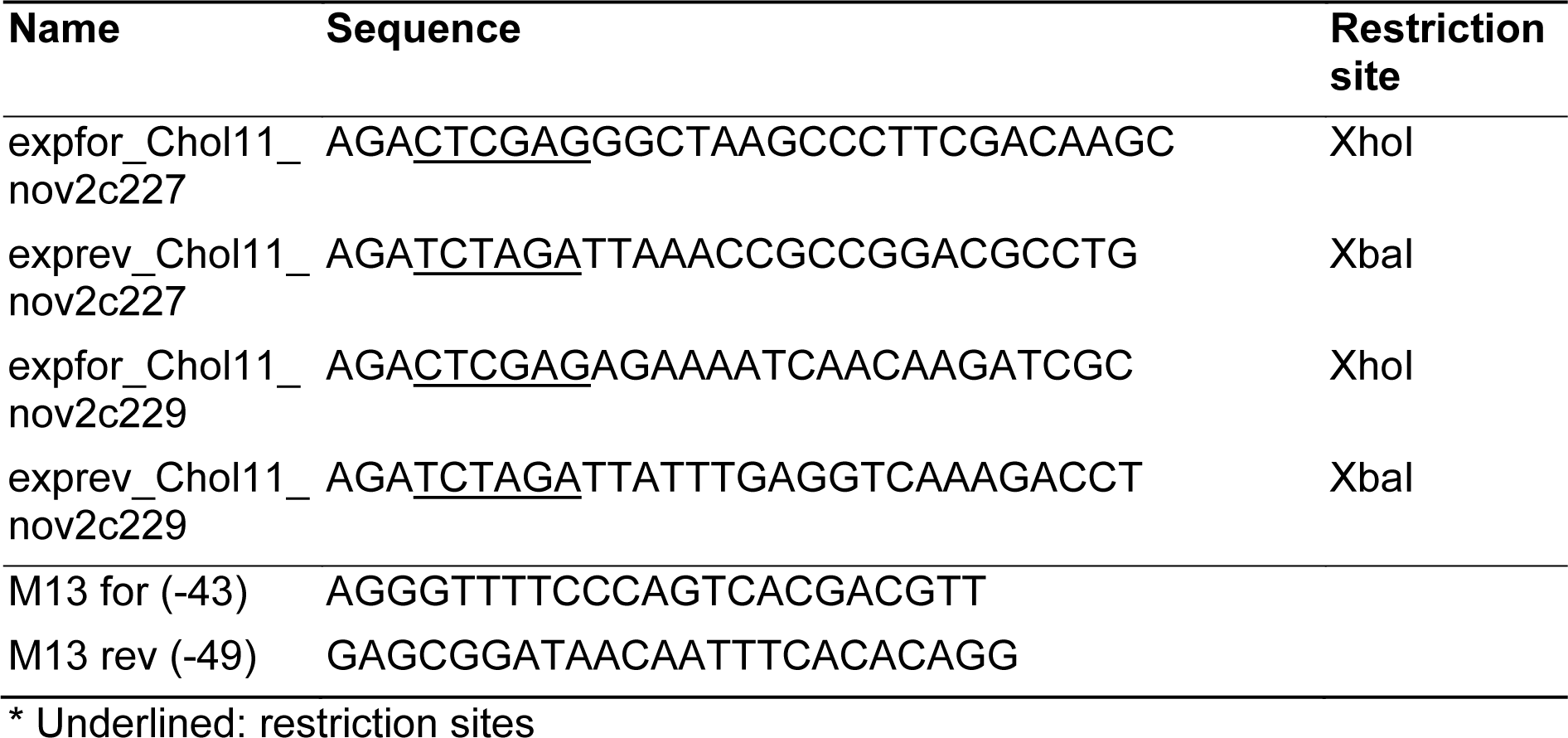
Primers used for construction of plasmids for heterologous expression.*

## Acknowledgements

The authors thank Karin Niermann and Kirsten Heuer (both Münster) as well as Christina Hinrichs (Oldenburg) for excellent experimental support and Florentin Schmidt for help with planning of cloning.

This work was funded by two grants of the Deutsche Forschungsgemeinschaft (DFG projects PH71/3-2 and INST 211/646-1 FUGG) to BP and a scholarship of the DAAD Stiftung in cooperation with the Prof. Dr. Bingel-Stiftung to FF.

